# Wide-field Calcium Imaging Reveals Widespread Changes in Cortical Connectivity Following Repetitive, Mild Traumatic Brain Injury in the Mouse

**DOI:** 10.1101/2022.02.22.481459

**Authors:** Samuel W. Cramer, Samuel P. Haley, Laurentiu S. Popa, Russell E. Carter, Earl Scott, Evelyn B. Flaherty, Judith Dominguez, Justin D. Aronson, Lukas Sabal, Daniel Surinach, Clark C. Chen, Suhasa B. Kodandaramaiah, Timothy J. Ebner

## Abstract

The physiologic basis underlying the long-term consequences of repetitive, mild traumatic brain injury (mTBI) remains poorly understood. Mild traumatic brain injury often results in brief loss of consciousness, impaired attention and concentration, memory problems, impulsivity, and headache, without objective findings on clinical imaging or examination. The effects of mTBI can persist and become cumulative with repetitive injury, suggesting global alterations in cortical networks. Using transparent polymer skulls, we performed mesoscopic Ca^2+^ imaging in mice to evaluate how repetitive mTBI alters patterns of neuronal interactions across the dorsal cerebral cortex. Spatial Independent Component Analysis (sICA) and Localized semi-Nonnegative Matrix Factorization (LocaNMF) were used to quantify changes in cerebral functional connectivity (FC). Repetitive, mild, controlled cortical impacts induce temporary neuroinflammatory responses, characterized by increased density of microglia exhibiting de-ramified morphology. These temporary neuro-inflammatory changes were not associated with compromised cognitive performance in the Barnes maze or motor function as assessed by rotarod. However, long-term alterations in functional connectivity were observed. Widespread, bilateral changes in FC occurred immediately following impact and persisted for up to 7 weeks, the duration of the experiment. Network alterations include decreases in global efficiency, clustering coefficient, and nodal strength, thereby disrupting functional interactions and information flow throughout the dorsal cerebral cortex. A subnetwork analysis shows the largest disruptions in FC were concentrated near the impact site. Therefore, repetitive mTBI induces a transient neuroinflammation, without alterations in cognitive or motor behavior, and a reorganized cortical network evidenced by the widespread, chronic alterations in cortical FC.

**Significance Statement:** More than 2.5 million individuals in the United States suffer minor traumatic brain injuries annually. Because these injuries are typically not associated with visible anatomic injuries or objective clinical findings, they were thought benign and fully recoverable. However, there is increasing awareness of the long-term deleterious consequences, particularly in patients who suffer repeated mTBI. Using long-term, mesoscopic neuronal Ca^2+^ imaging to characterize the dorsal cerebral cortical connectome following repetitive mTBI, we show extensive, persistent changes in functional connectivity, not only at the site of injury but throughout the cortex. These findings provide new insights into the pathophysiology of mTBI.

## Introduction

Mild traumatic brain injury (mTBI), defined as closed head injury resulting in a loss of consciousness and/or disorientation lasting < 30 minutes (Carroll et al., 2004a; Kristman et al., 2014), constitutes a major public health concern affecting at least 200-300/100,000 people per year (Lefevre-Dognin et al., 2021). Up to 60% of mTBI patients may show no evidence of brain injury on routine computerized tomography (CT) or conventional magnetic resonance imaging (MRI) (Yuh et al., 2013). Despite the absence of findings on medical imaging, mTBI is often accompanied by symptoms of nausea and vomiting, dizziness, headache, poor concentration, impaired memory, irritability, and emotional lability (Blennow et al., 2016). In addition to the immediate physical, cognitive, and behavioral symptoms, up to 30% of mTBI patients experience a post-concussive syndrome, characterized by CNS dysfunction > 3 months after injury (Blennow et al., 2016). Additionally, repeated mTBI leads to cumulative brain injury and poorer clinical outcome, including chronic traumatic encephalopathy (CTE) (Blennow et al., 2012; Smith et al., 2013). Given the prevalence and potential long-term adverse effects, it is imperative to understand the acute and secondary pathophysiological mechanisms of mTBI.

Characterizing the functional connectivity (FC) of spatially distributed and temporally correlated activity in regions throughout the brain as a “connectome” has emerged as a powerful tool for analyzing local and global network activity in the healthy and diseased brain (Biswal et al., 1995; Fornito et al., 2015; Caeyenberghs et al., 2017). Therefore, delineating FC after TBI provides an evaluation of injury-induced local and distributed changes, their association with cognitive and motor deficits, as well as neuroplastic changes over time. In moderate to severe TBI patients, both increased and decreased FC have been reported across distinct brain regions and functional networks (Rigon et al., 2016; Caeyenberghs et al., 2017; Palacios et al., 2017; Li et al., 2019), with hyper-connectivity the most common finding. In mTBI patients, decreases in FC have been noted when compared to healthy controls (Mayer et al., 2011; Johnson et al., 2012; Zhou et al., 2012). The complexity of the reported network changes after TBI is likely attributable to the heterogeneities in injury severity, distribution of injury (i.e., focal versus diffuse) as well as time from injury (Caeyenberghs et al., 2017). This inherent heterogeneity renders the study of mTBI in human subjects a fundamental challenge (Katz et al., 2015; Levin and Diaz-Arrastia, 2015).

Animal models offer advantages to study mTBI in the ability to precisely manipulate injury severity as well as the frequency and duration of observations after injury. Previous studies, however, are largely based on short term histological, immunochemical, and electron microscopy analyses at single, discrete time points after injury (Spain et al., 2010; Creed et al., 2011; Vascak et al., 2017, 2018). Lacking are studies of neuronal population and wide-field network dynamics, utilizing multi-time point assessments that are now possible with newer Ca^2+^ imaging technologies (for review see (Lin and Schnitzer, 2016; Cardin et al., 2020; Ren and Komiyama, 2021)).

The overarching goal of this study is to improve our fundamental understanding of mTBI using chronic wide-field Ca^2+^ optical imaging and network analysis tools to characterize local and global effects of mTBI on cerebral cortical activity in awake Thy1-GCaMP6 mice (Dana et al., 2014). An advantage of the wide-field Ca^2+^ imaging is that it provides a direct measure of neuronal activity in cortical excitatory neurons (Yizhar et al., 2011; Waters, 2020). Consistent with mTBI in human subjects, only transient changes in neuroinflammation followed mTBI, without deficits in working memory or motor function. However, we document persistent, widespread, bilateral changes in FC following repetitive cortical impacts, compared to sham controls.

## Material and Methods

### Fenestrated See-Shell design and fabrication

Transparent polymer skulls (i.e., See-Shells) were fabricated in a multistep process as described previously (Ghanbari et al., 2019; West et al., 2021). Briefly, See-Shell frames were 3D-printed out of polymethylmethacrylate (PMMA) (RSF2-GPBK-04, Formlabs Inc.). A 50 μm thick polyethylene terephthalate (PET) film (MELINEX 462, Dupont Inc.) was trimmed to match the See-Shell frame and bonded to it using epoxy adhesive (Scotch-Weld™ DP100 Plus Clear, 3M Inc.). Prior to implantation, a circular fenestration was created in the PET film using a hot solder iron tip at 550–600 °F to provide access for the tip of the controlled cortical impact (CCI) device. The fenestration was positioned over the right primary (M1) and secondary (M2) motor cortices (**Fig. 1A**).

**Figure 1.**
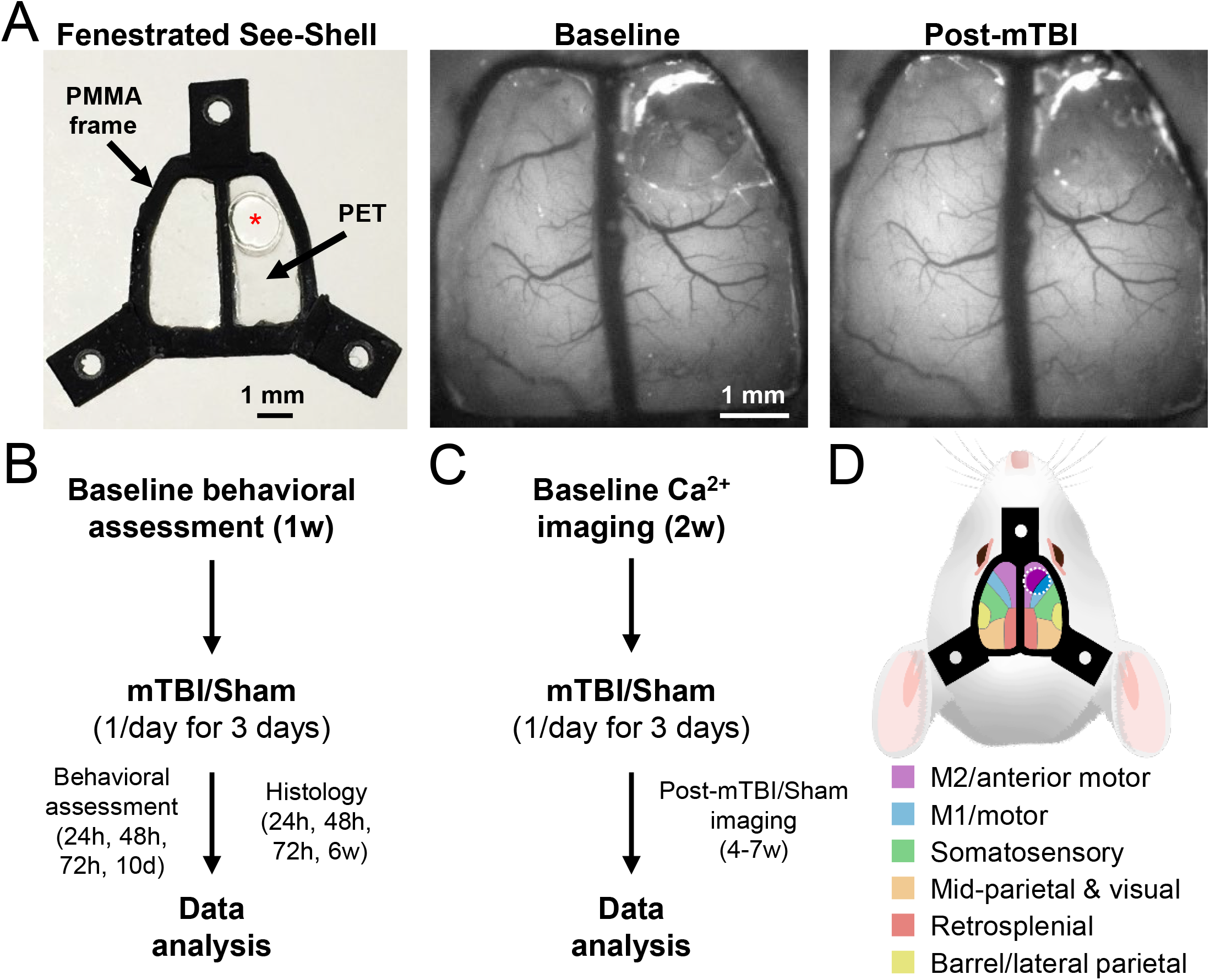
Implant and experimental design. **A**. (*left*) Image of the fenestrated See-Shell (* indicates opening in the PET). (*middle*) Baseline fluorescence image from a Thy1-GCaMP6f mouse 2 weeks after implantation of the fenestrated See-Shell. The PET opening is covered with a thin layer of a silicon polymer (Kwik-Sil). (*right*) Image from the same mouse 2 days post-mTBI. **B & C**. Experimental design of the behavioral and histological assessments (***B***) and the Ca^2+^ imaging (***C***). **D**. Illustration of the cortical regions (color coded to match legend) imaged through the fenestrated See-Shell. White-dashed circle denotes the opening in the PET.

### In vivo surgical implantation

All of the animal studies were approved by and conducted in conformity with the Institutional Animal Care and Use Committee of the University of Minnesota. We used Thy1-GCaMP6f mice (#024276, Jackson Laboratories) for Ca^2+^ imaging and C57BL/6 mice (#000664, Jackson Laboratories) for all other aspects. The procedure for implantation of the See-Shells was performed as described previously (Ghanbari et al., 2019). Mice were anesthetized with a mixture of 1–3% isoflurane and oxygen (0.6 mL/min). SR Buprenorphine (0.2 mg/kg, ZooPharm) and Meloxicam (2 mg/kg, Bayer Pharmaceutics) were administered subcutaneously for analgesia. The scalp was shaved, and mice were placed in a stereotactic frame. Their eyes were covered with ophthalmic eye ointment (Puralube, Dechra Veterinary Products) and their body temperature was feedback regulated. Anesthetic depth was assessed every 15 minutes. The scalp and periosteum were removed, and a dental drill was used to perform a craniectomy (leaving the dura intact).

The See-Shell was aligned to the craniectomy and placed over the exposed brain, providing optical access to a large portion of the dorsal cerebral cortex (**Fig. 1A** and **D**). The edges of the See-Shell were secured to the cranium via cyanoacrylate glue (VetBond, 3M Inc.). Dental cement (S380, C&B Metabond, Parkell Inc.) was used to fully secure the See-Shell and a custom titanium head-plate to the skull. The fenestration was sealed using silicone (KWIK-SIL, World Precision Instruments). After implantation, mice recovered on a heating pad until ambulatory, then returned to a clean home cage and were administered meloxicam once daily for 3 days postoperatively.

### Controlled cortical impact procedure

Controlled cortical impact (CCI) is a method for inducing a reproducible, graded brain injury (Xiong et al., 2013). However, the degree of injury and associated hemorrhagic tissue that results from commonly used parameters would interfere with Ca^2+^ optical imaging of brain activity. Also, as our goal was to model mTBI in humans with limited tissue damage, a modified CCI injury was devised based on a previously established approach (Chen et al., 2014) using a CCI tip (2 mm diameter with a rounded tip) printed out of flexible PMMA (Flexible 80A, Formlabs). Prior to the first impact, a single subcutaneous dose of SR buprenorphine was administered, as well as a dose of meloxicam before each impact. Using the modified tip attached to the CCI device (Hatteras Instruments, Model PCI3000), an impact to a 1 mm depth at 0.4 m/sec with an 85 ms dwell was delivered perpendicular to the dorsal surface of the brain once per day over 3 days (**Fig. 1B** and **C**). The protective silicone seal overlying the fenestration in the See-Shell was removed and replaced for each impact. The injury was applied consistently to the right M1/M2 motor regions. Sham animals underwent the same procedure, including anesthesia, analgesia administration, and replacement of the silicone seal, without application of the CCI.

### Histology

A subset of mice (n = 18 sham control and n = 21 mice post-mTBI) were fully anesthetized in 5% isoflurane and transcardially perfused with phosphate-buffered saline (PBS; CAT# P5493-1L, Sigma Aldrich) followed by 4% paraformaldehyde (PFA; CAT# P6148-500G, Sigma Aldrich). Brains were extracted and fixed in 4% PFA at 4 °C for one week. Two days prior to slicing, the brains were transferred to 30% sucrose solution for dehydration. To achieve hemisphere differentiation, a shallow incision was made along the lateral side of the cortex on the hemisphere contralateral to the impact. Coronal slices (50 μm) were cut using a manual sliding microtome. The slices were stored in PBS at 4 °C for up to one week prior to staining. Slices were then kept in PBS solution containing Triton X-100 and blocking agent (Normal Serum Block, CAT# 927502, Biolegend) for 2 h after which they were incubated in the solution containing: 1:1,000 primary antibody [goat Polyclonal anti-Iba-1 (NB100-1028SS, Novus Biologicals); rabbit Polyclonal anti-GFAP (PA5-16291, ThermoFisher Scientific); and mouse Monoclonal anti-NeuN (MAB377, Sigma-Aldrich)] for 24 h at 4 °C. Slices were washed and incubated in solution containing the secondary antibody (1:400) conjugated with fluorophores [donkey Polyclonal anti-goat Cy5 (705-175-147, Jackson ImmunoResearch Laboratories); donkey Polyclonal anti-rabbit Alexa Fluor Plus 488 (A32790, ThermoFisher Scientific); donkey Polyclonal anti-mouse Rhodamine Red-X (715-295-150, Jackson ImmunoResearch Laboratories)]. Slices were then washed in PBS and mounted using Invitrogen ProLong Diamond Antifade Mountant with DAPI (P36962, Invitrogen, ThermoFisher). Using the cortical incision made previously, all slices were mounted in the same orientation.

### Epifluorescence imaging acquisition and quantification of GFAP and Iba-1 immunofluorescence

In addition to determining overall Iba-1 levels, we also characterized microglial morphology. Brain sections immunostained for GFAP and Iba-1 were imaged with a Leica DM6000 B epifluorescence microscope with a 20X (0.5 NA) objective and tiled using Leica LAS X software. Regions of interest (ROIs) were outlined on each brain slice image, one on each hemisphere (Schindelin et al., 2012). Quantification of Iba-1 expressing cell density and global GFAP expression was performed using ImageJ (NIH). Iba-1 expressing cells were counted within each ROI to quantify cell density. Global GFAP expression was quantified by measuring the percentage of pixels above a preset threshold with each ROI.

### Confocal image acquisition and characterization of microglial morphology

Brain sections immunostained for Iba-1 (described above) were imaged with an Olympus FV1000 scanning laser confocal microscope with a 40X (1.3 NA) objective. Cy5 was excited with a 635 nm laser. Confocal stacks were taken at 1024 × 1024 pixels and a step size of 0.57 μm. Images were preprocessed with a custom ImageJ script and quantified with 3DMorph (York et al., 2018) to determine the ramification index and number of branch points. Images were taken in the primary motor cortex in both superficial cortical layers 2/3 and deep cortical layers 5/6. As there were no differences in the microglial morphology from the superficial and deep layers, the results were combined.

### Rotarod

The rotarod (Ugo Basile Mouse Rota-Rod #47600) apparatus consisted of 5 lanes on a rotating cylinder separated by large discs. The motorized cylinder rotated with linearly increasing acceleration from 5 to 50 RPM, over the span of 3 minutes. If the mouse spun around the cylinder twice, consecutively or not, or fell off during the test, the mouse was removed from the rotarod and the time of failure was recorded. The maximum time allowed for mice to stay on the rotarod was 3 minutes. The mice were given 3 consecutive training days on the rotarod consisting of 4 trials each day. A 10-minute rest period was applied in between trials. Mice were then tested weekly for 3 weeks. Rotarod evaluation before and after mTBI was performed on 12 mTBI and 13 sham mice.

### Barnes maze

The Barnes maze consisted of a 60 cm diameter circular table with 19 false holes and 1 escape hole all with a 5 cm diameter cut along the edge of the table. The false holes were cut at a depth of 1 cm into the table that did not go all the way through. The escape hole was cut through the table, allowing access to an escape chamber beneath. The maze was placed in a 2 × 2 × 2 m box draped with black curtains. Visual cues were placed on 3 of the walls. The arena was lit with four 60 W incandescent light bulbs mounted above the maze.

During the acquisition phase, the mice completed 4 trials per day for 4 consecutive days (Day −5 to −2 relative to CCI, see **Fig. 3B**). Animals were placed in a cylinder in the center of the maze in the dark to prevent visual orientation prior to the trial. To begin each trial, the lights were turned back on, the cylinder was lifted, and the mice were given 3 minutes to explore the maze and enter the escape hole. If the mouse did not enter the escape hole within 3 minutes, it was manually placed inside the hole for a period of 1 minute and then transferred back to its home cage. To eliminate olfactory cues, the maze and escape chamber were cleaned with 70% EtOH between trials. On each of these days, the location of the escape hole remained in the same position.

**Figure 2:**
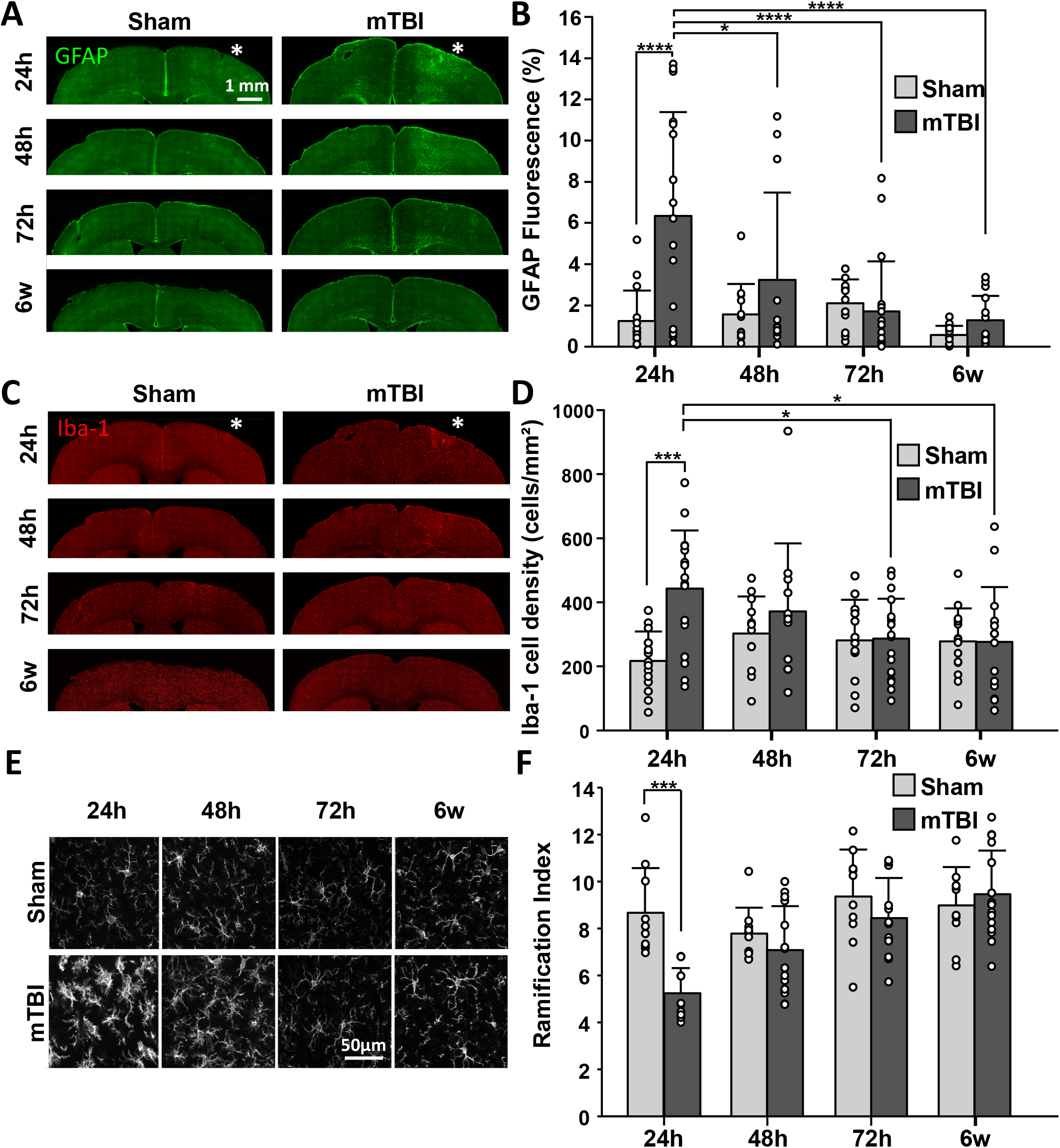
Changes in GFAP and Iba-1 expression following mTBI. **A & C**. Example immunostaining for GFAP (***A***) and Iba-1 (***C***) over time for sham (*left*) and mTBI (*right*) animals. * denotes ipsilateral site of the PET opening. **B & D**. Quantification of GFAP expression (***B***) and Iba-1 cell density (***D***). mTBI induced significantly higher global GFAP expression at 24h compared to sham animals (n=5/6, sham/mTBI; p < 0.0001), while it decreased significantly from 24h to 48h, 72h, and 6w following mTBI (n=6; p = 0.0195, p < 0.0001, p < 0.0001, respectively). Significantly higher Iba-1 cell density occurred at 24h in mTBI compared to sham (n=5/6, sham/mTBI; p < 0.0001), and decreased significantly from 24h to 72h and 6w for mTBI animals (n=6; p = 0.0121, p = 0.0121, respectively). **E**. Confocal images of Iba-1 showing microglia activation at 24h in mTBI (*bottom*) compared to sham animals (*top*). **F**. Microglial ramification was significantly lower in mTBI animals at 24h compared to sham (n=5/5 sham/mTBI, p < 0.05). Open circles in all plots are individual data points. *, ***,**** denotes p < 0.05, 0.001, and 0.0001, respectively.

**Figure 3:**
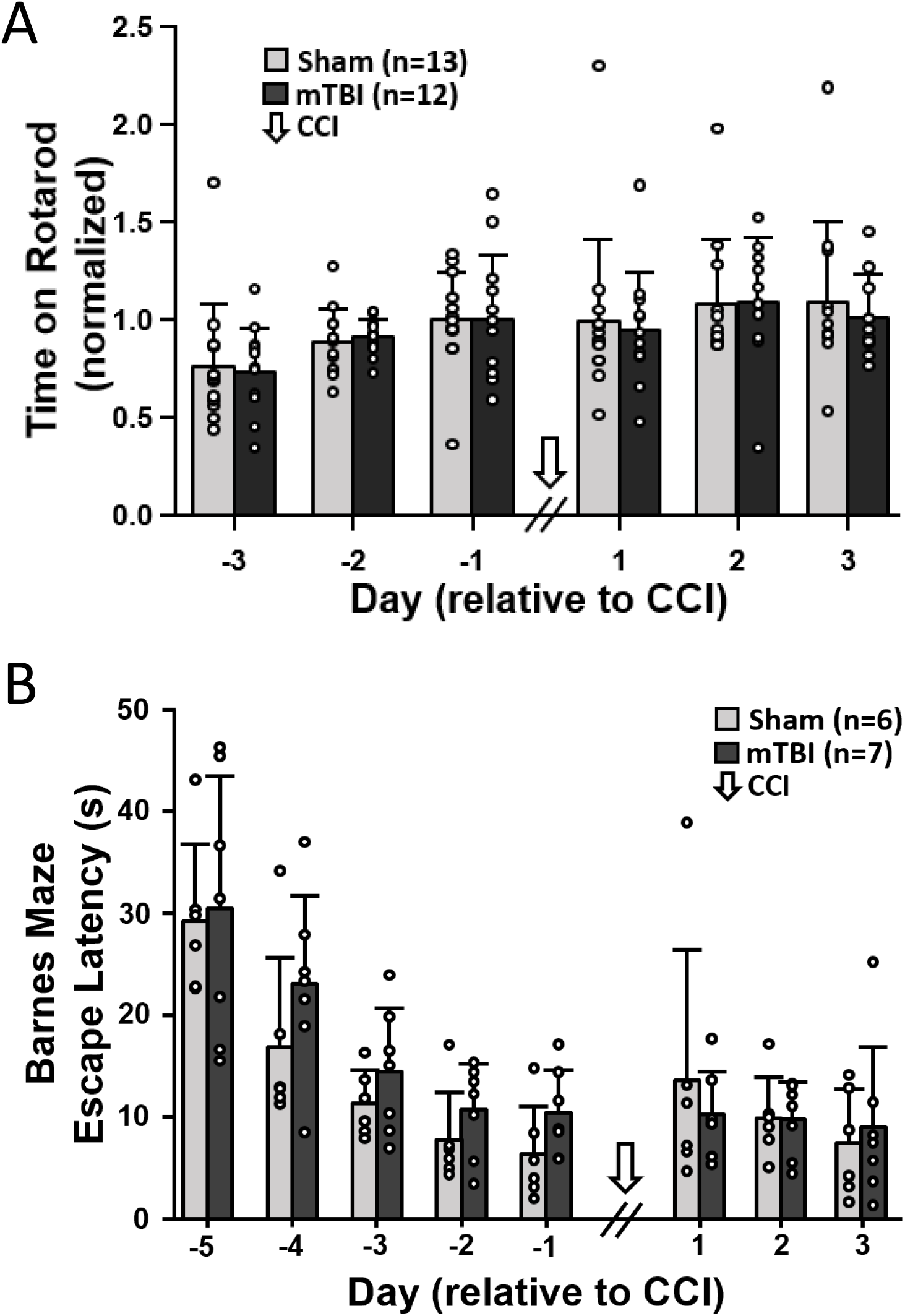
Motor and cognitive function are unaffected by mTBI. **A**. Rotarod performance between sham and mTBI animals. Data points presented as the average of 4 daily trials per animal, normalized to Day −1. **B**. Barnes maze escape latency between sham and mTBI animals. Latencies were averaged for the 4 daily trials per mouse, except probe days (Days −1 and 3) where only 1 trial was performed. White arrow denotes the time of CCI.

A probe trial was conducted on Day −1 relative to CCI. For this singular trial, the mouse was given 90 seconds to search the maze and enter the escape hole while in the same position as acquisition trials. The CCI protocol was administered over the next 3 days. The relearning phase was conducted on Days 1 and 2 post-CCI. During this phase, the table was rotated 135° and the same procedure as the acquisition phase was followed. Day 3 post-CCI consisted of a second probe trial where the maze remained in the same orientation as the relearning trials. Barnes maze evaluation before and after mTBI was performed on 7 mTBI and 6 sham mice.

### Barnes maze analysis

Videos of each trial were recorded with an overhead camera (BlackFly S USB3, Point Grey) and analyzed off-line for pose estimation with DeepLabCut (Mathis et al., 2018). By labeling the head, center of the body, and base of the tail, a custom MATLAB script tracked pose positions and trajectories to determine the latency to first interaction with the escape hole and number of hole poke errors. Escape latency was defined as the time from when the mouse was released to when the marker on the head initially crossed the outline of the escape hole. Poke errors were counted for each instance the point on the head crossed the outline of a false hole. Consecutive errors on the same hole were included if the delay was more than 75 ms between pokes.

### Wide-field Ca^2+^ imaging

Head-fixed mice were acclimatized on a low-friction, horizontal disk treadmill that allowed for natural movements and were imaged using a Nikon AZ-100 macroscope with a Plan Apo 1X objective. Single-photon fluorescence optical imaging was performed using a sCMOS camera (Andor Zyla 4.2, Oxford Instruments) controlled with MetaMorph (Molecular Devices Inc.). Neuronal activation is coupled with increased blood flow, driving light absorption with a peak ∼530 nm by the oxygenated blood, shortening the duration of the GCaMP fluorescence response (Ma et al., 2016). Therefore, dual-wavelength Ca^2+^ imaging was performed to mitigate the hemodynamic effects, as well as other Ca^2+^-independent fluorescence changes such as flavoprotein autofluorescence (Vanni and Murphy, 2014; Allen et al., 2017; Musall et al., 2019; Jacobs et al., 2020; MacDowell and Buschman, 2020). Ca^2+^-dependent (470 nm, blue light) and Ca^2+^-independent (405 nm, violet light) GCaMP6f signals were interleaved using a Cairn OptoLED driver (Cairn OptoLED, P1110/002/000; P1105/405/LED, P1105/470/LED). An excitation filter (ET480/40, Chroma) was placed in front of the 470 nm LED, then both light sources were combined into the parallel light path of the macroscope through a dichroic mirror (425 nm, Chroma T425lpxr), and then reflected off a second dichroic mirror (505 nm, Chroma T505pxl) to the brain. Cortical GCaMP6f emission passed back through the second dichroic into the camera. Exposure time for each frame was 18 ms, synchronized via TTL pulses from a Cambridge Electronics 1401 (Cambridge Electronic Design Limited) acquisition system that controlled both LEDs and the camera. Frames were captured at 40 Hz (20 Hz per channel) with 256 × 256 pixels per image.

Using the variable magnification function, the field-of-view was optimized to image the exposed dorsal cortical surface (6.2 mm x 6.2 mm) with a spatial resolution of 24.2 μm x 24.2 μm per pixel. During an imaging session, 6 to 10 trials (6000 frames, 5 min) were recorded with 1-5 min inter-trial interval. Mice studied with the Ca^2+^ imaging were subjected to the same CCI procedure described above, with 2 weeks of baseline recordings prior to, and up to 7 weeks post-CCI.

### Ca^2+^ imaging analysis using spatial Independent Component Analysis (sICA)

Fluorescence image preprocessing followed established protocols. Each trial was motion corrected using a subpixel image registration algorithm (Guizar-Sicairos et al., 2008). At the pixel level, both Ca^2+^-dependent and Ca^2+^-independent signals were detrended and then normalized to the mean fluorescence. The hemodynamic correction was performed by subtracting the normalized Ca^2+^-independent signal from the normalized Ca^2+^-dependent signal, equivalent to previously described in *Materials and Methods* (Ma et al., 2016; Jacobs et al., 2020; MacDowell and Buschman, 2020; West et al., 2021).

For each mouse, motion corrected data across all sessions were registered to a common frame to eliminate mounting-related variability using an affine transform function in MATLAB Then, a binary mask separating the brain from the background was determined for each mouse. All subsequent analysis was restricted to the pixels over the cerebral cortex. After registration, the hemodynamic corrected Ca^2+^ signal was denoised by removing, from each pixel over the brain, a common noise signal defined as the mean of all pixels over the background (not covering the brain) using linear regression.

Following denoising, imaging data from all the trials recorded were concatenated and compressed using single value decomposition (SVD). We kept the first 200 components that explain over 85% of the variability. From the SVD components, we computed 60 spatial independent components (ICs) and their time courses using the Joint Approximation Diagonalization of Eigenmatrices (JADE) algorithm that decomposes mixed signals into ICs by minimizing the mutual information with a series of Givens rotations (Cardoso, 1999; Makino et al., 2017; Sahonero-Alvarez, 2017). This method provides a blind segmentation of the cerebral cortex based only on statistical properties of the Ca^2+^ fluorescence signal, free from prior assumptions regarding the functional or anatomical organization. The ICs were z-scored and thresholded so that values below 2 were set to zero and ICs covering less than 400 pixels (∼0.23 mm^2^) were discarded as physiologically irrelevant. Any IC that included multiple discontinuous regions, for example, homotopic cortical regions, was separated into separate ICs for subsequent analyses. Further, upon visual inspection, we eliminated ICs that were not associated with cortical activity, such as edge or vascular artifacts (Makino et al., 2017; West et al., 2021).

Functional connectivity (FC) of the ICs extracted was determined using the Pearson correlations between the IC time courses over 1-minute, non-overlapping intervals. We used different analytical approaches to quantify the global mTBI effects on FC based on the ICs. One approach focused on the effects on the stronger IC interactions. The mean correlation values for each recording week, thresholded above 0.3, determined the FC among the ICs in each mouse (**Fig. 5**) and a Similarity Index among the weekly FC networks was determined using a 2-D Pearson correlation between adjacency matrices. We compared the week prior to mTBI (W-1) to each of the other weeks with an ANOVA with Bonferroni’s post-hoc correction (p < 0.05, GraphPad Prism GraphPad Software). We also quantified the global mTBI effects on the full FC network. We computed the global modularity and global efficiency for each 1-minute adjacency matrix (Rubinov and Sporns, 2010). To determine the effect of the mTBI, the values of each descriptor were grouped into weekly post-mTBI sub-populations and a baseline sub-population pooling the two pre-mTBI weeks. The post-mTBI weekly values were compared to the baseline values using an ANOVA with Bonferroni’s post-hoc correction (p < 0.05, GraphPad Prism GraphPad Software).

**Figure 4:**
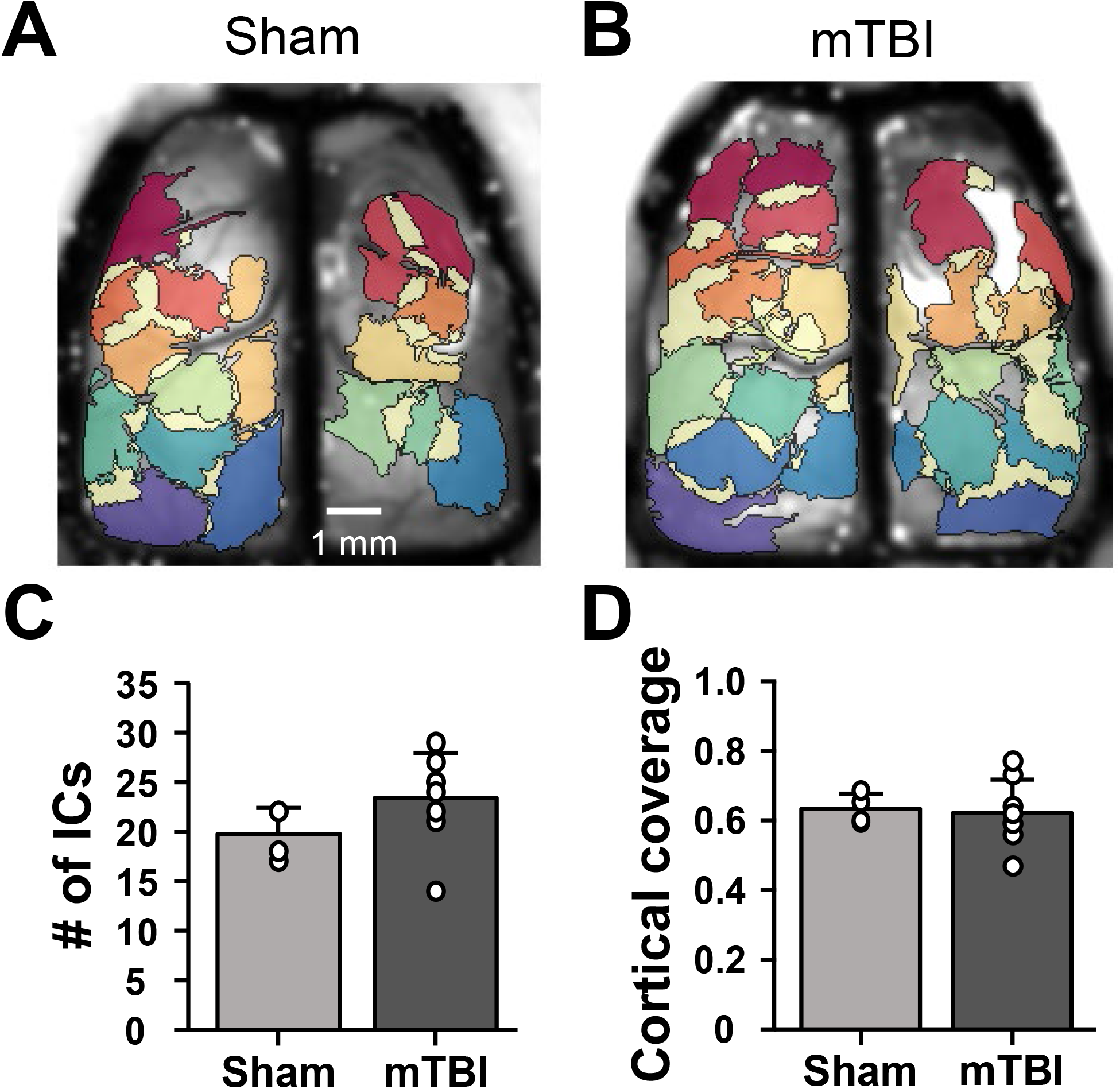
Functional cortical segmentation using sICA. **A and B**, Examples of cortical functional segmentation into ICs obtained over the entire optical recording data set, for a sham (**A**) and mTBI (**B**) mouse. **C**, Number of ICs did not differ between the sham and mTBI groups (t = −1.450, p = 0.178, Student t-test). **D**, Percent coverage by the ICs of the dorsal cortex, which did not differ between the sham and mTBI groups (t = 0.237, p = 0.817, Student t-test).

**Figure 5:**
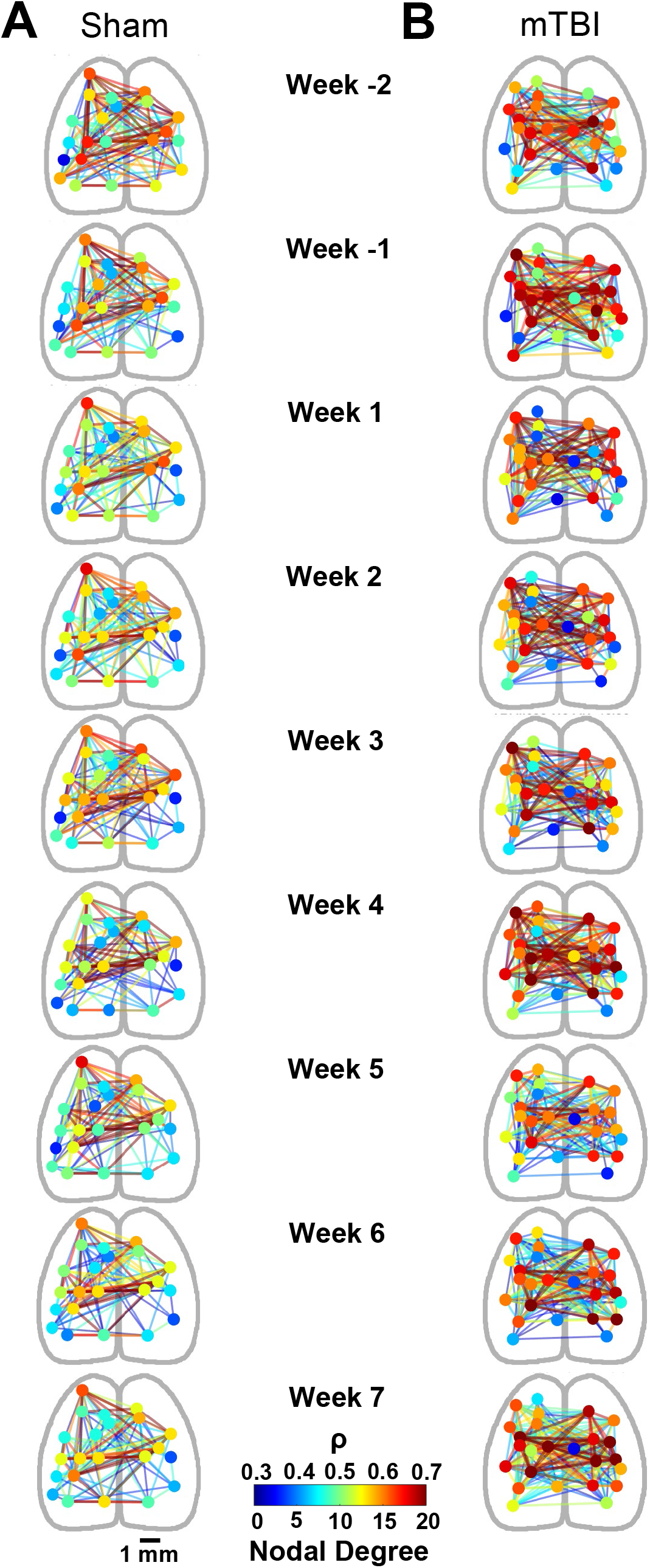
Functional connectivity networks. **A and B**. Examples of FC at each week in a sham (**A**) and a mTBI (**B**) animal. Node positions are the center of mass of ICs, plotted on a simplified outline of the dorsal cortex. The connections shown include only correlation coefficient (ρ) > 0.3. Nodes are color-coded to nodal degree. Edges are color-coded to ρ.

The second method evaluated the spatial structure of the larger changes of the network FC, due to mTBI, referred to as the Δ network. For each pair of ICs, we determined whether the pre-mTBI (baseline) and the post-mTBI correlation populations (grouped by week) came from different distributions. We used a Kolmogorov-Smirnoff test with Bonferroni correction to account for all IC pairs and an effect size threshold that required the difference between mean baseline (defined as the average of the two weeks pre-mTBI) and weekly mean to exceed 0.3. The IC pairs that met both criteria formed the weekly Δ network, describing the extent of the mTBI effect on cortical FC. At a group level, we used graph theory descriptors including nodal degree, nodal strength, and node betweenness (Rubinov and Sporns, 2010) to quantify the mTBI effects described by the Δ networks each week. We performed a 2-way ANOVA with weekly and mouse groups factorization on each descriptor population to determine differences between mTBI and sham groups.

### Ca^2+^ imaging analysis using localized semi-nonnegative matrix factorization (LocaNMF)

To provide additional insights into cerebral cortex functioning following mTBI, we performed an independent analysis using LocaNMF (Saxena et al., 2020). While sICA has many advantages to assess changes in functional connectivity (Calhoun and de Lacy, 2017), averaging across animals is challenging, as the number and position of the ICs varies across animals. LocaNMF uses the Allen Brain Atlas Common Cortical Framework (CCF) for the anatomical segmentation for all mice, facilitating averaging results across animals (Lein et al., 2007). LocaNMF is a form of multinomial principal component analyses, as opposed to blind source separation by sICA, providing an independent evaluation.

The masked, aligned, and hemodynamic corrected data generated from the preprocessing for the sICA was also analyzed following the procedures in (Saxena et al., 2020). Briefly, in the LocaNMF preprocessing, the data from each mouse was segmented into 1-minute time bins, then denoised and aligned to the CCF. LocaNMF was then run on each 1-minute time segment, allowing for 3 spatial components per atlas region with 100 minimum pixels required for a spatial component (as opposed to 400 used in sICA due to the size limitations of some of the atlas regions), 80% localization threshold, and capturing 99% of the variance in the data captured. The canonical correlation coefficients were calculated for each atlas region pair to generate the adjacency matrices over all 1-minute non-overlapping time segments. Only atlas regions that were visible after masking across all mice were used for subsequent analysis. Graph theory metrics of global efficiency, nodal strength, and clustering coefficient were calculated on each of the non-thresholded, weighted adjacency matrices. The distribution of each metric from each week post-mTBI or sham was compared to the distribution of the baseline period (the two weeks pre-mTBI or sham) by a 2-way ANOVA with Bonferroni’s post-hoc correction (p < 0.05).

A total of 25 atlas regions were found in common across all mice. These regions include: MOp: primary motor cortex; MOs: secondary motor cortex; RSP: retrosplenial cortex (agl: lateral agranular part; d: dorsal part); SSp: primary somatosensory cortex (bfd: barrel field; ll: lower limb; tr: trunk; ul: upper limb; un: unassigned); VIS: Visual cortex (al: anterolateral; am: anteromedial; p: primary; pm: posteromedial; rl: rostrolateral). Left (contralateral to impact) and right (ipsilateral to impact) hemispheres are denoted as _L and _R, respectively.

### Experimental design and statistical analysis

In addition to the FC statistical analyses described above, statistical evaluation of changes in immunofluorescence, microglial morphology, and behavioral measures between mTBI and sham animals were performed using an ANOVA with Bonferroni’s post-hoc correction (p < 0.05) using MATLAB (The MathWorks) or GraphPad Prism (GraphPad Software). In the text and figures, all values are reported as mean ± SD. Individual data points are plotted as open circles in all bar graphs, with the exception of **Figures 6, 8**, and **11** where the large number of points obscures the presentation. When describing the results of an experiment, “n” refers to the number of animals used.

**Figure 6:**
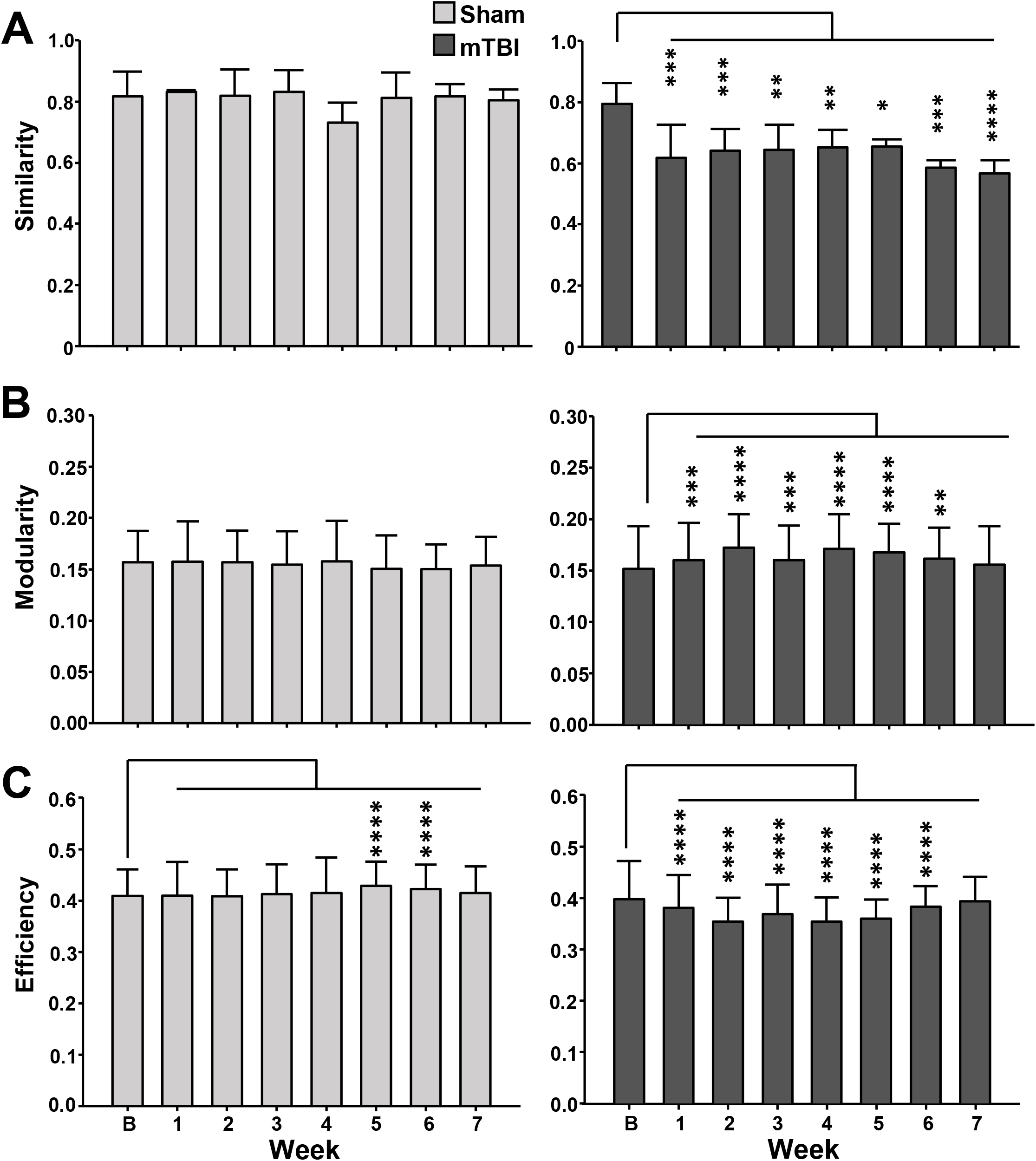
Functional connectivity network quantitative descriptors from sICA. **A**. Group summary plots of the Similarity Index. **B**. Group summary plots of the full network global modularity. **C**. Group summary plots of the full network global efficiency. All comparisons are relative to the group baseline (Week B). *, **,***,**** denotes p < 0.05, 0.01, 0.001, and 0.0001, respectively.

**Figure 7:**
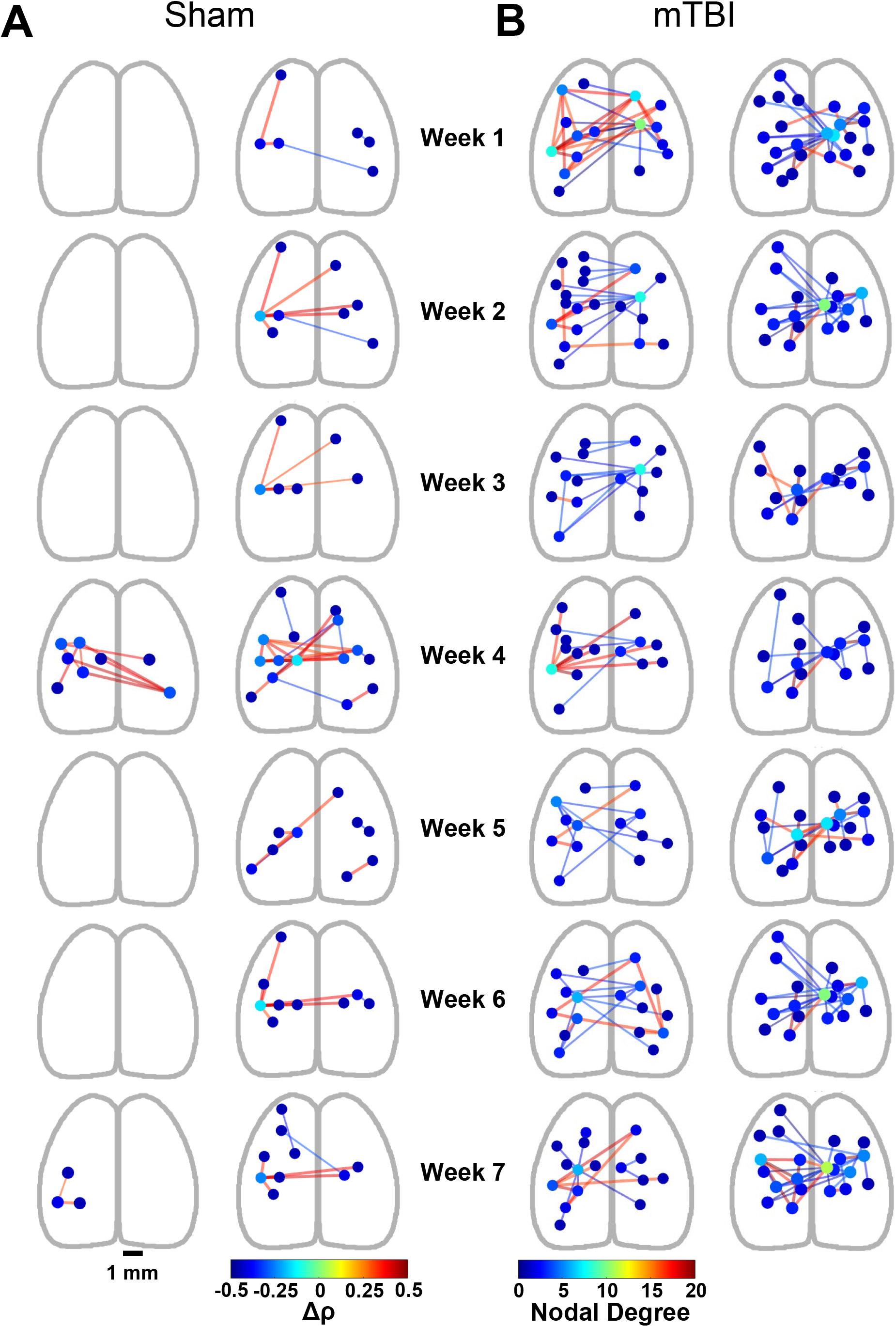
Δ networks. **A and B**. Example of the Δ networks at each week in two sham animals (***A***) and two mTBI animals (***B***). Nodes are color-coded to nodal degree. Edges are color-coded to change in correlation coefficient (Δρ, blue = decrease; red = increase).

**Figure 8:**
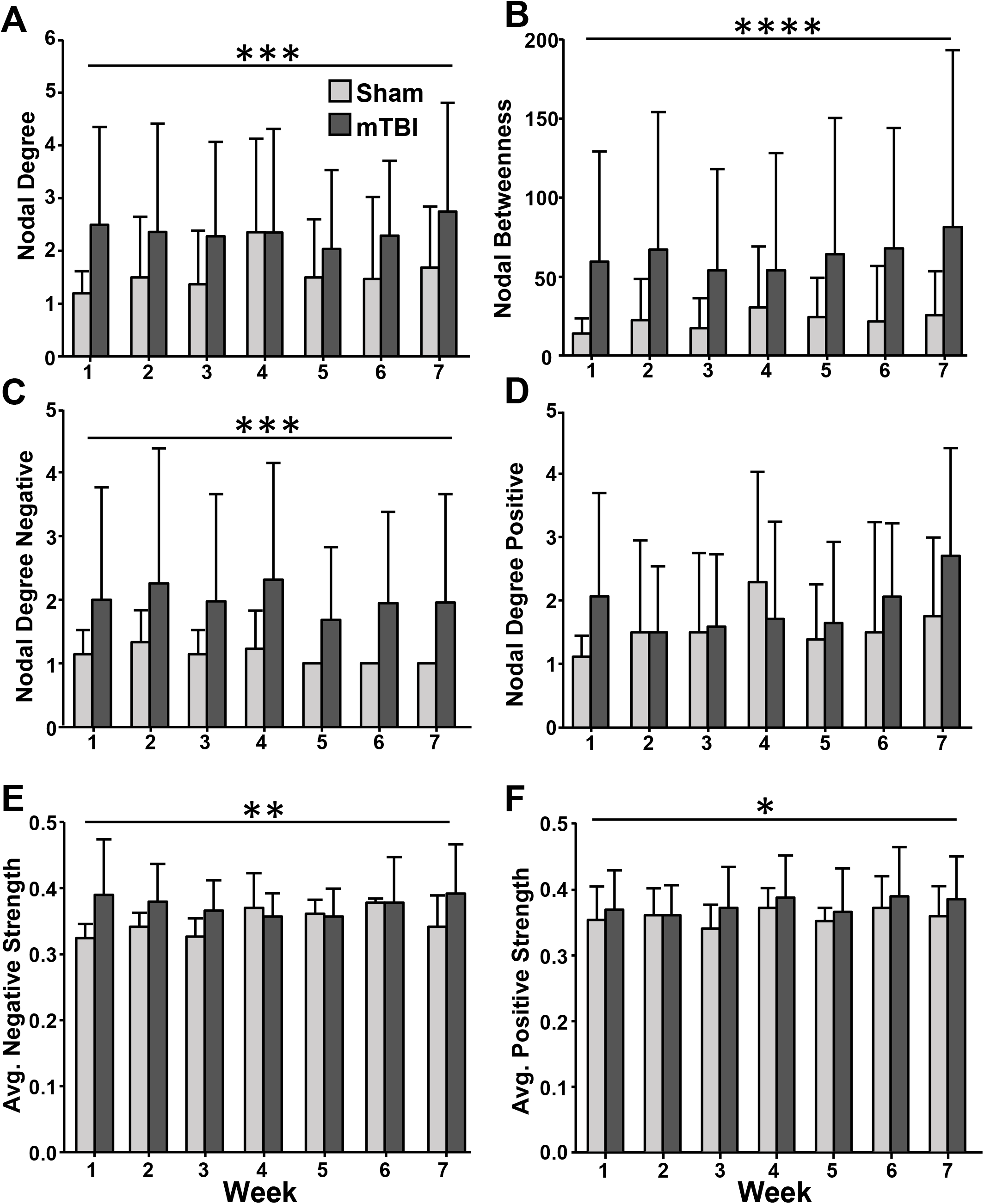
Group Δ network descriptors. **A**. Nodal Degree. **B**. Nodal Betweenness. **C**. Nodal Degree Negative – includes only correlation decreases (blue edges in Fig.6). **D**. Nodal Degree Positive – includes only correlation increases (red edges in Fig. 6). **E**. Average Negative Strength – mean of the correlation decrease magnitudes at each node. **F**. Average Positive Strength – mean of the correlation increases at each node. *, **,***,**** denotes p < 0.05, 0.01, 0.001 and 0.0001, respectively.

## Results

### Histological characterization of the mTBI model

Astrogliosis is a hallmark of a brain injury, including TBI (Burda et al., 2016). Therefore, to evaluate the extent of neuroinflammation induced by our mTBI model, histological evaluation of the astrocytic marker, GFAP, in brain sections from mTBI and sham animals was performed at 24, 48, and 72 h as well as 6 w post-injury (**Fig. 2A** and **B**). A significant increase in GFAP expression was observed at 24 h following mTBI compared to sham in the cortex ipsilateral to the injury (F(1,107) = 11.88, p = 0.0008). This increase in GFAP expression progressively decreased each day following injury and normalized by 72 h post-mTBI.

Neuroinflammation often correlates with clinical outcomes following TBI (Loane et al., 2014; Kumar et al., 2016; Loane and Kumar, 2016) and activated microglia are a critical element of this response. We examined microglial activation, via immunostaining for Iba-1 and found a significant increase in microglia cell density between mTBI and sham groups in the cortex ipsilateral to the injury (F(1,107) = 7.195, p = 0.00085, **Fig. 2C** and **D**). At subsequent time-points post-injury, the density of Iba-1 expressing cells normalized between mTBI and sham in the ipsilateral cortex. Therefore, the neuroinflammatory response evoked by our mTBI model was transient and normalized within 72 h post-mTBI. Finally, we did not observe any injury to subcortical structures such as the hippocampus or striatum, suggesting the injury was highly localized to the cortex.

To further characterize microglia activation, we quantified the 3-dimensional (3D) morphology of Iba-1 expressing cells. High-resolution confocal images show less ramified morphology and shorter branch length in microglia following mTBI at 24 and 48 h, characteristic of an activated state ((Loane et al., 2014; Loane and Kumar, 2016), **Fig. 2E**). Quantification of microglia morphology reveals a significant reduction in ramification 24 h following mTBI compared to sham (F(1,31) = 4.98, p = 0.0330), with no significant change at later time points (**Fig. 2F**). Taken together, our novel mTBI model exhibits transient neuroinflammation, in the form of astrogliosis and microglial activation, that subsides within 48-72 h after the injury.

### Motor and cognitive characterization of the mTBI model

To examine whether our mTBI model was associated with impaired motor function, rotarod testing was undertaken in a separate cohort of mTBI (n = 7) and sham (n = 6) mice. We observed no difference in rotarod performance between mTBI or sham animals (F(1,132) = 0.19, p = 0.6602; **Fig. 3A**), indicating that no overt motor deficit occurred following mTBI.

Barnes maze is a widely used spatial memory task which examines working memory. As no motor deficits were observed in the rotarod task, we expected any deficit in the ability to perform the task following mTBI would be due to impaired cognition (i.e, working memory). Across days, we found no significant difference in the latency to first interaction with the escape hole between sham and mTBI (F(1,88) = 1.94, p = 0.17; n = 6/7, sham/mTBI, **Fig. 3B**). Additionally, there was no significant difference in the number of poke errors prior to reaching the escape hole between the mTBI and sham groups (F(1,88) = 1.51, p = 0.22; n = 6/7, sham/mTBI). To test whether a cognitive deficit occurred past the short testing period above (3 days), a second group of mice was tested in the Barnes maze 10 days after injury. Despite the longer time following injury, there was no significant difference between mTBI and sham groups in either first latency (F(1,32) = 0.18, p = 0.67; n = 3/3, sham/mTBI) or poke errors (F(1,32) = 0.017, p = 0.90; n = 3/3, sham/mTBI), suggesting there are no cognitive impairments following our mTBI procedure. Taken together, the rotarod and Barnes maze results do not demonstrate deficits in motor function or working memory following mTBI, which is consistent with the paucity of overt clinical deficits in most humans within 1-3 months following mild TBI (Carroll et al., 2004b).

### Ca^2+^ imaging and functional connectivity of the dorsal cerebral cortex based on sICA

Wide-field Ca^2+^ imaging sessions were performed each week on additional mice (sham = 5, mTBI = 8) for 2 weeks prior to mTBI, followed by either 4 weeks or 7 weeks after mTBI (see **Fig. 1C**). Mice were head-fixed over a disk treadmill that allowed for spontaneous locomotion and wide-field Ca^2+^ imaging performed through the implanted polymer optical window (see *Materials and Methods*). After preprocessing and hemodynamic correction, each cortical surface was functionally segmented using sICA using the entire data set, including both pre and post mTBI recordings (420,000 to 650,000 frames per mouse). This resulted in a set of ICs and their associated time courses for each mouse that were used for the subsequent functional analyses.

The sICA segmentation of the dorsal cerebral cortex resulted in 18-22 ICs per mouse, consistent with previous reports (Makino et al., 2017; West et al., 2021). The ICs provide for a functional segmentation that covers the majority of the dorsal cerebral cortex in the imaging field for both sham and mTBI mice (**Fig. 4A** and **B**, respectively). Neither the number of ICs (t(10) = 1.45, p = 0.1776) nor the average coverage achieved (t(10) = 0.51, p = 0.6183) differed in the two groups **(Fig. 4C** and **D**). In the upper right quadrant, near the fenestration and impact site, ICs were identified in the mTBI mice as well as the sham, showing that the Ca^2+^ signals near the mTBI site were not completely disrupted. While there are similarities in the ICs, the exact location and size of the ICs differs for each mouse, as observed previously (West et al., 2021).

The FC was determined for each week, pre and post mTBI. The Pearson correlations between all pairs of IC time courses restricted to the week of interest were used to generate the FC adjacency matrix for each mouse. The FC network was plotted on an outline of the cortex, with the centers of mass of each IC as the network nodes coordinates and the edges as the strength of correlation, ρ, between nodes (only edges with ρ ≥ 0.3 are shown, **Fig. 5A** and **B**). As shown for an example sham mouse, the network is qualitatively stable over 9 weeks (**Fig. 5A**).

To quantify the stability of the IC-based network, we calculated a Similarity Index (see *Materials and Methods*), comparing the week prior to mTBI (W-1) with each of the other weeks. For the sham group, the Similarity Index is above 0.75 for all comparisons and there are no significant differences across weeks (F (7, 21) = 0.97, p = 0.48, **Fig. 6A**). In contrast, mTBI induces changes in the network structure, resulting in significant decreases of the Similarity Index for weeks 1 to 7 (F(7, 44) = 5.94, p < 0.0001, **Fig. 6A**). Analysis of the full network shows that global modularity in the sham group presents no significant changes relative to baseline (F(7,1909) = 1.819, p = 0.0795, **Fig. 6B**), while the mTBI group presents significant increases in weeks 1 to 6 (F(7,3449) = 22.00, p < 0.0001, **Fig. 6B**). The global efficiency of the full network in the sham group presents an increase in weeks 5 and 6 (F(7,1909) = 10.66, p < 0.0001, **Fig. 6C**) relative to baseline, as opposed to decreases in weeks 1 to 6 for the mTBI group (F(7,3449) = 127.5, p < 0.0001, **Fig. 6C**). Together, the decreased Similarity Index and increase modularity following mTBI indicate the network is more divided, and the decreased global efficiency suggests a reduced ability to integrate neural activity through the network.

The complete networks are complex as are the differences in the global network metrics, so to better understand the larger FC changes, we determined the networks that include only the node pairs with the largest significant changes in correlation (Δ networks, see *Materials and Methods)*. To determine the significant changes in correlations we computed the FC over 1-minute non-overlapping intervals for each week and compared each node pair’s weekly correlation population to a baseline population that combined the 2 weeks before mTBI (see *Materials and Methods*). In addition to a Bonferroni corrected Kolmogotov-Smirnoff test to determine the significant differences between baseline and the post-CCI/sham groups, we also required that the absolute difference between the mean correlations exceed 0.3. Examples from the sham group reveal remarkably sparse Δ networks (**Fig. 7A**), with only a small number of nodes and edges differing from baseline. For mice in the mTBI group, the Δ networks are extensive, covering large expanses of the cortical area. Many nodes and edges change relative to baseline, with the changes dominated by decreases in correlation strength (**Fig. 7B**). The connectivity changes are not restricted to the cortical impact site in the right M1/M2 motor cortices. Instead, network changes are present throughout the dorsal cerebral cortex, both ipsilateral and contralateral to the site of impact, and the changes persist for up to 7 weeks following mTBI.

To further quantify the changes induced by mTBI, we assessed node betweenness, nodal degree (positive and negative), and average nodal strength (positive and negative), of the Δ networks, comparing the sham and mTBI groups. Nodal degree (F(1,714) = 20.89, p < 0.0001) and node betweenness (F(1,710) = 33.14, p < 0.0001) are significantly greater in mTBI than in sham mice (**Fig. 8A** and **B**), reflecting the increased complexity of the Δ networks in the mTBI group relative to sparse Δ networks in sham animals (see also **Fig. 6**). Further, based on the sign of the correlation differential, we dissected the Δ networks into nodes with increased correlation strength (positive) and nodes of decreased strengths (negative). Following mTBI, nodal degree negative increases relative to sham (F(1,439) = 11.58, p = 0.0007, **Fig. 8C**) while the nodal degree positive in the mTBI group remains similar to the sham group (F(1,391) = 3.62, p = 0.059, **Fig. 8D**). Also following mTBI, both average negative strength (F(1, 439) = 7.02, p = 0.0084, **Fig. 8E**) and positive strength (F(1, 391) = 6.29, p = 0.0126, **Fig. 8F**) are significantly larger relative to the sham group, showing that at both positive and negative nodes in the Δ network correlation strength changes. However, the change in positive strength is small with only select nodes exhibiting increased correlation strength (see **Fig. 7B**). Therefore, the Δ networks following mTBI are dominated by pairs of nodes with large decreases in correlation strength. For all Δ network measures, neither the week nor interaction (group by week) terms were significant and implies stability of the Δ networks over the time period evaluated.

To better understand the changes in the spatial structure across the cortex following mTBI, we plotted the location of the Δ network nodes on the Allen Brain CCF (**Fig. 9**), combining the weekly data from the sham and mTBI groups. For each CCF region, we computed for the Δ networks the density of nodes in each area, normalized by the area of the region (in mm^2^) and number of animals. First, these population maps illustrate the sparseness of the Δ network in sham mice versus the greater density of changed nodes throughout the dorsal cerebral cortex in mTBI mice. Second, the maps highlight that the largest disruptions in the network are concentrated in and most often involve ICs in the right motor cortex, the site of the impact. Regions in the contralateral, homotopic cortex also show a high density of change following mTBI. Interestingly, there is the suggestion of a delayed effect in the contralateral motor cortex illustrated by an increase in the change density in weeks 6 and 7.

**Figure 9:**
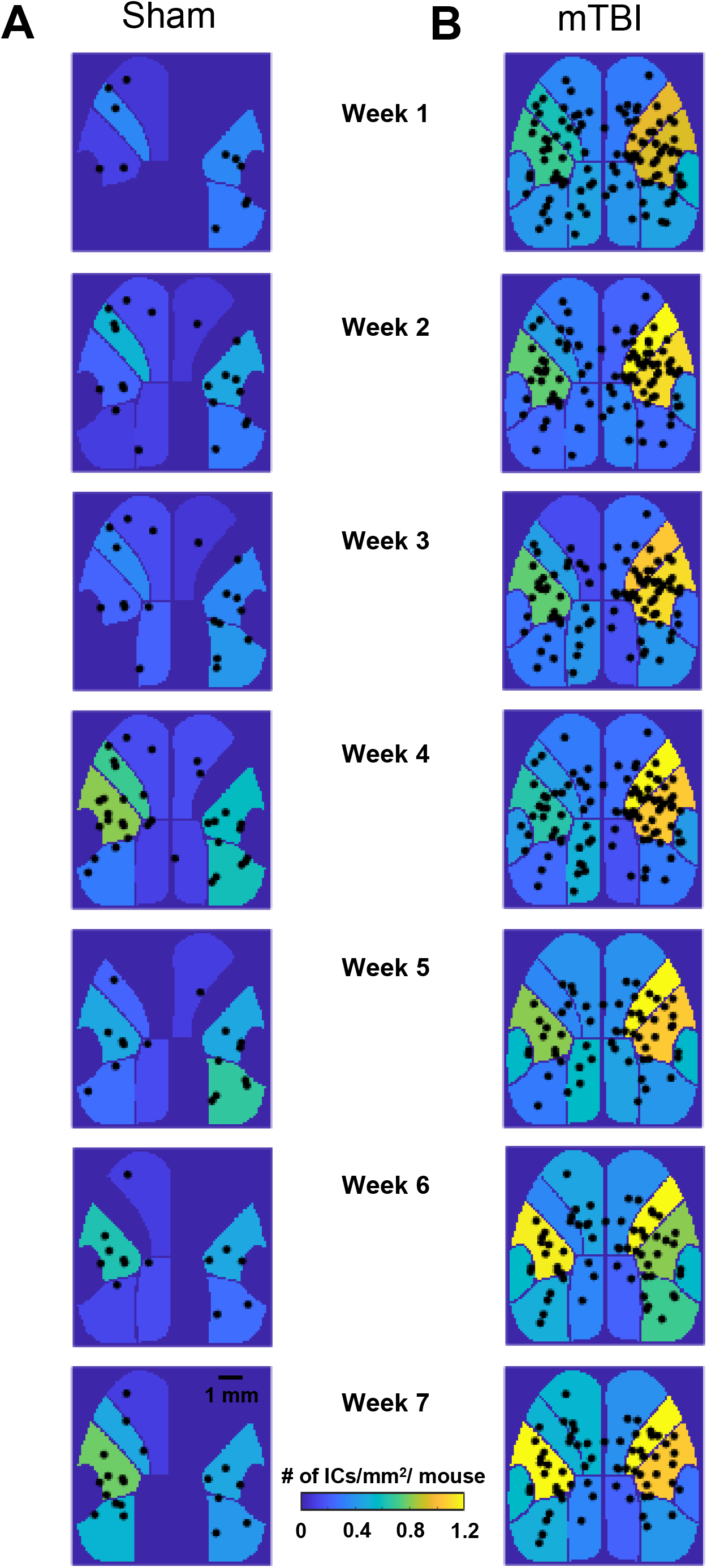
Effects of mTBI on IC density. **A and B**. Center of mass of ICs that showed significant changes following sham (***A***) or mTBI (***B***), plotted on a canonical segmentation of the cerebral cortex (Allen Brain Atlas). Each cortical region (see **Fig. 1D** for labels) is color-coded to the density of ICs normalized to the number of mice in each group and week.

### Functional connectivity of the dorsal cerebral cortex based on LocaNMF

We undertook a second analysis of the changes in FC using LocaNMF (Saxena et al., 2020). In addition to providing an independent evaluation of how mTBI alters cerebral cortical interactions, LocaNMF allows for averaging across mice, as the anatomical segmentation is based on the CCF (see *Materials and Methods*). Examination of the average adjacency matrices based on the canonical correlations confirmed the overall stability of the network in sham mice across the entire time span (**Fig. 10A**), as observed with sICA. Several subnetworks with high correlations among the component subregions, including motor, retrosplenial, and visual regions are evident. Following mTBI, the adjacency matrices show an overall decrease in connectivity, particularly for these motor, retrosplenial, and visual networks (**Fig. 10B**). Over the 7 weeks following mTBI, global efficiency significantly decreased (F(7,5755) = 19.23, p < 0.0001, **Fig. 10C**), confirming the reduced network integration observed with sICA.

**Figure 10.**
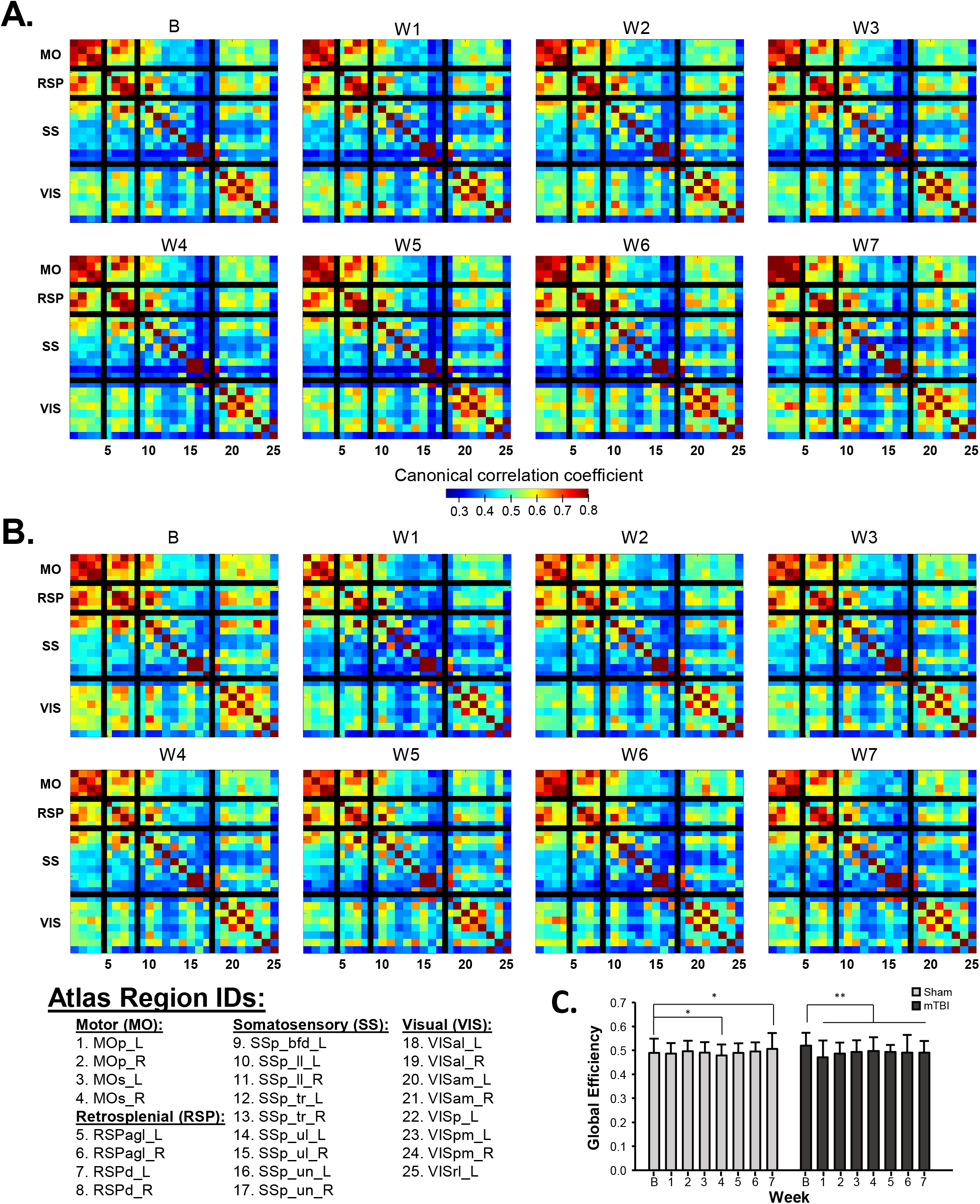
Large-scale decreases in FC following mTBI revealed by LocaNMF. **A and B**. Canonical correlation matrices from all sham (***A***) and mTBI (***B***) mice over time across the Allen Brain Atlas regions in all mice. Regions are grouped into generic functional areas which are separated by black lines. Atlas region abbreviations are provided (for full names, see *Material and Methods*). **C**. Global efficiency calculated from the weighted correlation matrices from each 1-minute imaging time bin. * and ** denotes p<0.05 and 0.0001, respectively.

The analysis of network statistics at the nodal level focused on nodal strength (**Fig. 11A**) and the clustering coefficient (**Fig. 11B**). For visualization, we plotted the average magnitude of the significant changes relative to baseline for each cortical region, for the sham and mTBI cohorts. For both nodal degree and the clustering coefficient, there are widespread, persistent, bilateral decreases following mTBI (**Fig. 11A** and **B**, right columns). The largest amplitude decreases were ipsilateral to the injury site and within the motor, retrosplenial, and somatosensory subnetworks. The sham cohort (**Fig. 11A** and **B**, left columns) shows fewer significant changes of smaller magnitude. Also, significant changes in the sham cohort were a mixture of spatially isolated increases and decreases, as opposed to the global decreases seen in the mTBI cohort.

**Figure 11.**
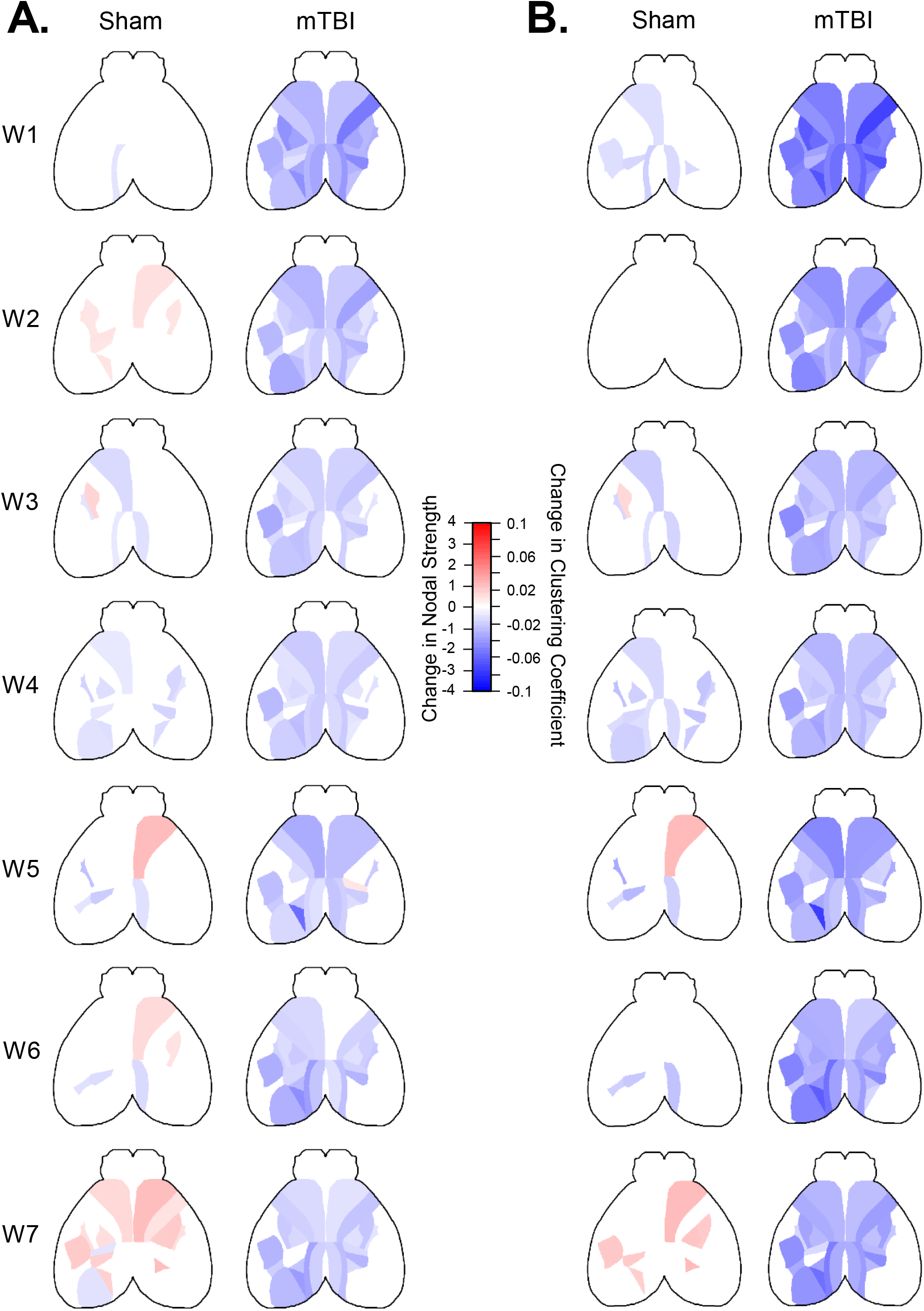
Significant decreases in functional connectivity metrics following mTBI revealed by LocaNMF. **(A)** Changes in nodal strength from sham (*left*) and mTBI (*right*) color-coded to magnitude of the change for each atlas region that was significantly different from baseline. (**B)** Same as in (***A*)** but for the clustering coefficient. Only atlas regions that remained significant following Bonferroni’s post-hoc correction (p < 0.05) are shown.

## Discussion

We report a new murine model to investigate the effects of repetitive mTBI on dorsal cerebral cortical networks using mesoscopic Ca^2+^ imaging of excitatory neurons. We followed cerebral cortical neuronal activity for up to 7 weeks post-mTBI in awake animals to determine the time-course of both acute and subacute changes of cerebral cortical FC. The fundamental findings are that both local and global cerebral cortical network alterations occur following repetitive mTBI, as well as brief, transient increases in local neuroinflammation without appreciable motor or cognitive deficits.

We show that mTBI acutely perturbed the cerebral cortical network observed during baseline, resulting in a decrease in network similarity. Moreover, the decrease persisted for 6 weeks, suggesting a long-term reorganization of the cortical network. Both sICA and LocaNMF reveal decreased global efficiency following mTBI, indicative of a reduction in functional integration with loss of parallel information transfer (Achard and Bullmore, 2007; Rubinov and Sporns, 2010). The decrease in nodal strength further highlights that mTBI reduces overall functional connectivity. The clustering coefficient is a topological measure of the relationship between any pair of neighbors for each node that provides an assessment of the capacity for information integration within this triad of nodes. While not a direct measure of spatial segregation, the decreases in the clustering coefficient show that mTBI alters the potential for functional specialization by decreasing the number of small, local, functional cortical motifs in the network (Rubinov and Sporns, 2010). Further, the decreases in nodal strength and clustering coefficient are widespread, involving most of the imaged cortical regions.

The extent of the changes was mapped by the Δ networks, defined by the IC derived nodes with the largest changes in correlation. The Δ networks following mTBI have increased complexity compared to the sham subjects, reflected in increased nodal degree and node betweenness. Both increases and decreases occur, requiring partition into Δ networks of positive and negative sub-networks. Following mTBI, nodal degree is significantly higher in the negative but not the positive subnetwork, showing that the changes in the cortical connectivity are largely determined by correlation decreases. While there was an increase in nodal density in the Δ networks throughout the cortex, the highest density occurred in somatomotor areas near the impact site. The LocaNMF results also find that these regions exhibit some of the largest decreases in FC.

Although very different mathematically, sICA and LocaNMF provided mutually re-enforcing results. Making no prior assumptions, sICA extracts the cortical regions within each mouse whose neural activity is maximally statistically independent from each other. While a powerful approach to investigate changes within each mouse, sICA poses a challenge to incorporate data across mice. By restricting the localization of activity to the CCF, LocaNMF enables generating cortical networks with identical node locations. This allows averaging FC across mice and achieve a more complete description of the network changes and their cortical locations.

### Comparison to human studies

While some human subject studies describe decreases as well as increases in FC following mTBI (Johnson et al., 2012; Iraji et al., 2016; Churchill et al., 2017, 2018; Palacios et al., 2017; Champagne et al., 2019), persistent disruptions in FC in multiple sensory and cognitive networks are reported in mTBI patients that develop the post-concussive syndrome (Stevens et al., 2012; Vakhtin et al., 2013). Moreover, such reduced connectivity persists for up to 4 months (Johnson et al., 2012; Shumskaya et al., 2012; Zhou et al., 2012; Nordin et al., 2016) Taken together, these results suggest that mTBI in patients is commonly characterized by a significant reduction in connectivity. Similarly, both sICA and LocaNMF analyses in our study show that the dominant FC change is a decrease. The FC findings, when combined with the mild limited histopathology and behavioral findings, imply that our model recapitulates human disease.

When compared to moderate or severe injury, the effects of mTBI on cortical networks appear to be fundamentally different. Most human FC studies report hyper-connectivity for moderate and severe TBI (Hillary et al., 2011; Sharp et al., 2011; Palacios et al., 2013; Bernier et al., 2017; Mohamed et al., 2021). Mild and severe TBI result in structural lesions, a critical difference from mTBI that likely contributes to the hyper-connectivity.

### Wide-field Ca^2+^ imaging provides a novel measure of mTBI on FC

To our knowledge, this is the first report of wide-field Ca^2+^ imaging before and after mTBI, providing new insights into the reduced FC. Imaging based on fMRI or similar approaches are indirect measures of neuronal activity. Our hemodynamic-corrected imaging provides a direct measure of neuronal activity and demonstrates that the reduced FC is due to decreased neuronal activity/function. The Ca^2+^ fluorescence signals detected in the mice (Thy1-GCaMP6f) using our single-photon imaging are primarily from the excitatory neuronal activity in layers II/III (Yizhar et al., 2011; Ma et al., 2016; Waters, 2020). Cortical layers II/III provide intrahemispheric projections throughout the cortex and interhemispheric projection via callosal axons to equivalent locations in the contralateral cortex (Zhang et al., 2016; Chovsepian et al., 2017). The reduction in FC observed is also consistent with the reduction in synchrony in beta band connectivity observed in patients following mTBI (Rier et al., 2021). Therefore, the present methodology complements other measures of connectivity by more precisely reflecting neuronal specific changes and their contribution to FC. While our study investigated the effects on FC following mTBI to the motor cortex, it could easily be adapted to examine how mTBI in other brain regions affect FC.

### Comparison to previous rodent models of mTBI

In addition to the chronic wide-field Ca^2+^, our injury model differs from previous rodent mTBI models using closed head impact (for review see (Bodnar et al., 2019)) as well craniotomy dependent CCI used in the present study (for review see (Siebold et al., 2018)). Few experimental studies have longitudinally characterized changes in cerebral cortical networks at multiple times in the same animal following mTBI beyond one or two time points (Harris et al., 2016; Meningher et al., 2020; Yang et al., 2021). Furthermore, MRI is typically acquired from anesthetized animals. Common anesthesia protocols alter neurovascular coupling (Pan et al., 2015), hemodynamic activity (Ku and Choi, 2012), the extent and magnitude of correlated BOLD response (Bajic et al., 2017), and FC (Mandino et al., 2019; Reimann and Niendorf, 2020).

Several previous studies used moderate/severe TBI that resulted in anatomic lesions and focal neurological deficits (Harris et al., 2016; Yang et al., 2021). As our goal was to study mTBI in awake animals with no/minimal cortical pathology and no hemorrhage, we used a 2 mm, flexible PMMA tip on the impactor as well as very conservative impact parameters (Chen et al., 2014). Compared to previous studies (Siebold et al., 2018), our CCI parameters (1 mm depth, 0.4 m/sec) would be classified as “mild”. However, 0.4 m/sec is approximately an order of magnitude lower than most previous mTBI studies using CCI. Several studies classified as mild that used higher speeds resulted in greater tissue damage at the injury site as well as parenchymal damage in deeper structures, including the hippocampus, as well as behavior deficits (for review see (Siebold et al., 2018)). The minimal and transient histopathological changes, similarities in the number of ICs extracted and lack of overt behavioral findings in our study are consistent with the impact parameters utilized. In fact, several mTBI studies using more severe impact parameters failed to demonstrate gross motor deficits (Sauerbeck et al., 2012; Bolte et al., 2020) or impairment in spatial memory (Hemerka et al., 2012).

Previous FC studies in rodents used MRI methodologies. Two rat studies using CCI injury (Harris et al., 2016; Yang et al., 2021) and resting state fMRI, reported increases in connectivity following injury compared to the present results. However, the CCI produced significant cortical damage in both studies, even in animals classified as having mild injury (Harris et al., 2016). Therefore, these results are consistent with the hyper-connectivity in humans following moderate/severe TBI. A mouse study using diffusion MRI to examine structural connectivity following a closed head impact (Meningher et al., 2020) reported decreased global efficiency and clustering early that returned to normal as well as changes to both hemispheres, as observed here with Ca^2+^ imaging.

### Final comments

Our results, when taken together with previous human and experimental studies, highlight that repetitive mTBI can lead to persistent changes in cortical FC. The acute perturbation of cortical connectivity may be due to neural dysfunction during the transient tissue inflammation. In turn, this perturbed network responds to the injury under constraints of maintaining brain functionality. The result is a reorganized cortical network that maintains gross functionality, though in a suboptimal manner. These findings bear relevance to symptoms experienced by many mTBI patients.

## Acknowledgements

We would like to thank Lijuan Zhuo for assistance with animal surgeries. We thank Alexander Cramer at the University of Minnesota University Imaging Center (SCR_20997) for assistance in generating graphics and 3D printing, and the Mouse Behavior Core for behavioral testing. The work was supported in part by NIH R61/R33 NS115089, NIH R01 NS111028, Minnesota SCI-TBI fund (Grant Contracts: 143722 and 191546), and University of Minnesota’s MnDRIVE (Minnesota’s Discovery, Research, and Innovation Economy) initiative.

## Notes

***Conflict of interest:*** The authors declare no conflicts of interest pertaining to this work.

### Competing Interest Statement

The authors have declared no competing interest.

